# Genome assemblies for seven families of birds from the Global South

**DOI:** 10.1101/2025.07.31.667860

**Authors:** K. L. Vinay, Naman Goyal, Ashwin Warudkar, Chiti Arvind, V. V. Robin

**Affiliations:** Department of Biological Sciences, Louisiana State University, Baton Rouge, LA, USA; Department of Biology, Indian Institute of Science Education and Research Tirupati, Tirupati, Andhra Pradesh, India

**Keywords:** The Global South, Old World Tropics, Western Ghats, Reference genomes, Birds

## Abstract

Tropical regions are biodiversity-rich, yet remain underrepresented in the availability of genomic resources, as is evident in the Western Ghats of India, a biodiversity hotspot with high endemism. Here, we present high-quality, de novo genome assemblies for seven birds, representing seven families distributed in the Western Ghats: Black-naped Monarch (Monarchidae: *Hypothymis azurea*), Indian Yellow Tit (Paridae: *Machlolophus aplonotus*), Brown-cheeked Fulvetta (Leiothrichidae: *Alcippe poioicephala*), Malabar Trogon (Trogonidae: *Harpactes fasciatus*), Blue-bearded Bee-eater (Meropidae: *Nyctyornis athertoni*), Malabar Whistling-Thrush (Muscicapidae: *Myophonus horsfieldii*), Orange-headed Thrush (Turdidae: *Geokichla citrina*). Using a hybrid Oxford Nanopore long reads – Illumina short reads approach, we assembled genomes with sizes ranging from 1.03 to 1.13 Gbp. All assemblies demonstrated high contiguity and completeness (BUSCO scores >97%, UCEs >4799). Repeat masking identified ∼10% of the genomes as interspersed repeats, and functional annotations yielded an average of 9,619 protein-coding genes per species. Comparative analysis showed our assemblies had significantly higher contiguity than the median of existing avian genomes on NCBI (Wilcoxson test, p = 0.00226). Our genome assemblies fill a key geographic and taxonomic gap in the genomic data and provide a foundational resource for evolutionary and ecological research in the Old-World tropics.

## 1. Introduction

The Global South - encompassing the regions of Latin America, Africa, and much of Asia - harbors extraordinary biological diversity and yet remains underrepresented in large-scale genomics studies (Coelho et al. 2023; Reddy 2014). Historically, the imbalance in research funding and focus has meant that scientific endeavors have concentrated heavily in the Global North (Soares et al. 2023; Liverpool 2021). This creates a critical data gap; while genomic references for numerous temperate species continue proliferating, tropical and subtropical taxa are often left without genome assembly (Linck and Cadena 2024). Consequently, key questions about adaptation, evolutionary history, and conservation remain unanswered for many of the most biodiverse regions (Hortal et al. 2015). Addressing this disparity is essential to forming a truly global understanding of biodiversity and evolutionary processes shaping diversity hotspots such as the Western Ghats of India.

The Western Ghats of India represent one of the world’s most critical biodiversity hotspots and are recognized for the remarkable array of endemic flora and fauna (Shanavas, Sumesh, and Haris 2016). Spanning over 1,600 kilometers along the western edge of peninsular India, this mountain chain harbors vast tracts of tropical and subtropical moist broadleaf forests, montane grasslands, and montane forests (Singh and Chaturvedi 2017; Robin and Nandini 2012). With significant topographical complexity and a wide range of microclimates, the Western Ghats sustain a high level of endemism across multiple taxonomic groups (Goyal et al. 2025; Bharti et al. 2021; Ansari and Gill 2016; Dahanukar, Raut, and Bhat 2004). For birds, in particular, the Western Ghats provide habitats that range from lowland rainforest to high elevation peaks, each with a unique biogeographical gradient (Raman, Joshi, and Sukumar 2005; Raman 2006; Pramod et al. 1997; Robin and Nandini 2012). As a result, several bird lineages have diverged within the Western Ghats, resulting in distinct species and subspecies (Robin et al. 2017; Ramachandran et al. 2017; Wickramasinghe et al. 2017). Despite the clear ecological and evolutionary significance, the Western Ghats remain underrepresented in studies incorporating genomic and Sanger datasets (Reddy 2014). Birds have long served as a model system in evolutionary biology due to their diverse life history strategies, pronounced morphological variations, and well-documented fossil and observational records (Brusatte, O’Connor, and Jarvis 2015). Yet, compared to global bird studies, many avian lineages in South Asia remain understudied from a genomic perspective and lack whole-genomic resources. This gap is especially conspicuous within the Western Ghats, where the endemism is high.

High-quality genome assemblies are indispensable for modern biological research. They provide the foundational blueprints for a wide range of studies, from comparative genomics and phylogenetics to functional and conservation genetics (Rhie et al. 2021; Zadesenets, Ershov, and Rubtsov 2017; Whibley, Kelley, and Narum 2021). High-quality genome assemblies allow researchers to identify genes involved in adaptation (Wang et al. 2023), characterize structural rearrangements and repetitive elements (Flynn et al. 2024), and understand the genetic basis of behavior, such as migration (Weissensteiner et al. 2024) and elucidate patterns of genome organization (Edwards et al. 2025). In contrast, studies reliant on fragmented assemblies often fail to capture the full spectrum of genomic variation, limiting the power to understand the evolutionary process (Whibley, Kelley, and Narum 2021; Rhie et al. 2021). Recent advances in long-read sequencing have opened up many opportunities for assembling non-model organism genomes from telomere to telomere (Li and Durbin 2024; Garg et al. 2024; Makova et al. 2024). Oxford Nanopore sequencing technology offers rapid, high-throughput long-read sequencing data spanning several thousand kilobases to megabases, allowing the generation of high-quality genome assemblies.

Here, we present the first de novo assemblies and functional annotations for seven bird species –Black-naped Monarch (Monarchidae: *Hypothymis azurea;* BNMO), Indian Yellow Tit (Paridae: *Machlolophus aplonotus;* IYTI), Brown-cheeked Fulvetta (Leiothrichidae: *Alcippe poioicephala;* BCFU), Malabar Trogon (Trogonidae: *Harpactes fasciatus;* MATR), Blue-bearded Bee-eater (Meropidae: *Nyctyornis athertoni;* BBBE), Malabar Whistling-Thrush (Muscicapidae: *Myophonus horsfieldii;* MWTH), Orange-headed Thrush (Turdidae: *Geokichla citrina;* OHTH) from seven families originating from Western Ghats of India (Figure 1) using a hybrid Oxford Nanopore – Illumina approach. We also compare our de novo assemblies to the existing avian genome assemblies in the National Center for Biotechnology Information (NCBI).

**Figure 1:**
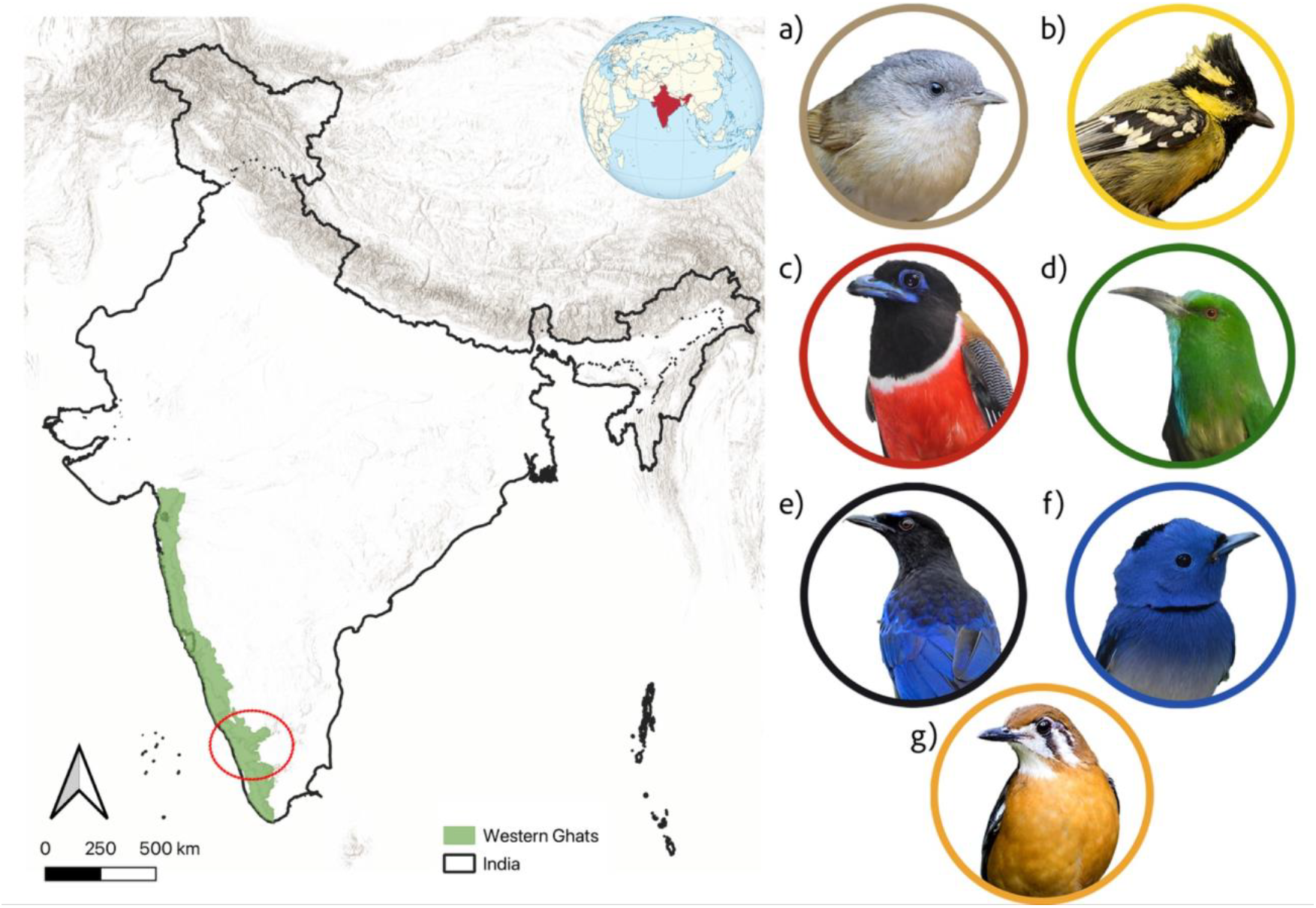
Sampling location and representative images of the seven study species of birds from the Western Ghats. The left panel shows the extent of Western Ghats highlighted in green on the Indian map and the colored dotted circle on the map corresponds to the Shola Sky Islands - the sampling locations for all species The right panel is the representative photographs – a) Brown-cheeked Fulvetta (*Alcippe poioicephala;* BCFU), b) Indian Yellow Tit (*Machlolophus aplonotus;* IYTI), c) Malabar Trogon (*Harpactes fasciatus;* MATR), d) Blue-bearded Bee-eater (*Nyctyornis athertoni;* BBBE), e) Malabar Whistling-Thrush (*Myophonus horsfieldii;* MWTH), f) Black-naped Monarch (*Hypothymis azurea;* BNMO), g) Orange-headed Thrush (*Geokichla citrina;* OHTH). All photographs are from Wikimedia Commons under the CC BY-SA license.

## 2. Materials and Methods

### 2.1 Sample Collection

We conducted fieldwork across the Western Ghats of India and collected blood samples from the brachial vein following Robin, Sinha, and Ramakrishnan 2010. We stored the blood in Queen’s Lysis buffer (QLB) (Seutin, White, and Boag 1991) and froze it till DNA extraction.

### 2.2 DNA Extraction and Qualitative Assessment

Genomic DNA was extracted from frozen blood using Qiagen DNeasy Tissue and Blood kit (GmbH, Germany) following the manufacturer’s protocol with modifications: i) an increased amount of Proteinase K, and ii) a more extended incubation period. Isolated gDNA quality was assessed using a 1% agarose gel, and concentration was determined using a Qubit 4.0 fluorometer. gDNA was then sent to a commercial sequencing facility for library preparation and sequencing. For long reads, we targeted approximately 10x coverage on Oxford Nanopore (ONT hereafter) PromethIon (R10.4.1 Flowcell) using Dorado-v7.3.11 in high accuracy mode for basecalling. For short reads, we targeted approximately 30x coverage on Illumina NovaSeq 6000 for paired-end reads.

### 2.3 Read Pre-processing

The quality of ONT reads was assessed using NanoPlot v1.43.0 (De Coster and Rademakers 2023), and any residual adapters were trimmed via Porechop v0.2.3 (Wick et al. 2017; De Coster and Rademakers 2023). Adapter-free reads were further filtered to remove low-quality (q>7) bases using Chopper v0.7.0 (De Coster and Rademakers 2023). For Illumina reads, we first used Fastp (Chen et al. 2018) to assess the quality, and then, using Trimmomatic v0.39 (Bolger, Lohse, and Usadel 2014), we removed the adapters and low-quality bases. Adapter-trimmed paired-end short reads were used to generate the K-mer frequencies (with K=20) using Jellyfish v2.2.6 (Marçais and Kingsford 2011), and GenomeScope2 (Ranallo-Benavidez, Jaron, and Schatz 2020) was then used to determine the genome size and heterozygosity for each of the samples (See Supplementary Figure S1)

### 2.4 Nuclear Genome Assembly

Quality filtered and adapter trimmed ONT reads were de novo assembled using Flye v2.9.5-b1801 (−- nano-raw) (Kolmogorov et al. 2019) with the corresponding genome size. We then used FCS-gx and FCS-adapter v0.5.4 (Astashyn et al. 2024) to screen the ‘draft’ assemblies for contamination; if found, they were removed before proceeding further. We removed any contigs corresponding to mitogenomes, identified using de novo assembled mitogenomes (see below for details) and minimap2 v2.28-r1209 (Li 2018). We then polished the assemblies using both long-read and short-read-based strategies. First, Medaka v2.0.1 (recommended by Nanopore Tech) was used with long reads to polish with the dna_r10.4.1_e8.2_400bps_hac@v4.3.0:consensus model in the ‘inference’ module, followed by one round of polishing with Racon v1.4.0 (Vaser et al. 2017) using long reads. Subsequently, with short reads, two rounds of POLCA (distributed under MaSuRCA v4.1.2, Zimin and Salzberg 2020) were applied to obtain the ‘polished’ assemblies. Redundant haplotigs were removed using purge_haplotigs v1.1.3 (Roach, Schmidt, and Borneman 2018)with a coverage cut-off -l 2 -m 8 (& -m 10 for *Machlolophus aplonotus*) -h 190 (See supplementary Figure S2). We then scaffolded the assemblies using ntLinks v1.3.11 (Coombe et al. 2021, 2023) with ntLinks_rounds (w=250, k=32, rounds=5) with gap-filling. Since ntLinks gap-filling uses raw reads, polishing the assembly post-gap-filling is recommended. So, we ran a final round of polishing with POLCA and short reads to obtain the final assemblies. We used assembly-stats v1.0.1 (https://github.com/sanger-pathogens/assembly-stats) and compleasm v0.2.6 (Huang and Li 2023) with the Benchmarking Universal Single-Copy Orthologs (BUSCOs) aves_odb10 dataset to assess the assembly quality and completeness. We also harvested the Ultra Conserved Elements (UCEs) following the standard method (https://phyluce.readthedocs.io/en/latest/tutorials/tutorial-3.html) with Phyluce 1.7.3 (Faircloth 2016) as a quality metric to determine the completeness of the assembly. Final scaffolds were renamed using a custom Python script.

### 2.5 Mitochondrial Genome Assembly

We first used findMitoReference.py from the MitoHiFi suite (Uliano-Silva et al. 2023) to find the available closest sister species mitogenome (listed in supplementary Table S1), which was then used as a seed and the reference for NOVOPlasty v4.3.5 (Dierckxsens, Mardulyn, and Smits 2016) to de novo assemble the mitogenomes using short reads. For two species that failed the mitogenome assembly via NOVOPlasty, we used mitohifi.py (Uliano-Silva et al. 2023) with the scaffolded nuclear assembly as an input to assemble the mitogenomes. Obtained contigs were annotated using MitoAnnotator v4.09 (Zhu et al. 2023; Ato et al. 2018; Iwasaki et al. 2013). Refer to Figure 2 for a schematic methodology of the assembly steps.

**Figure 2:**
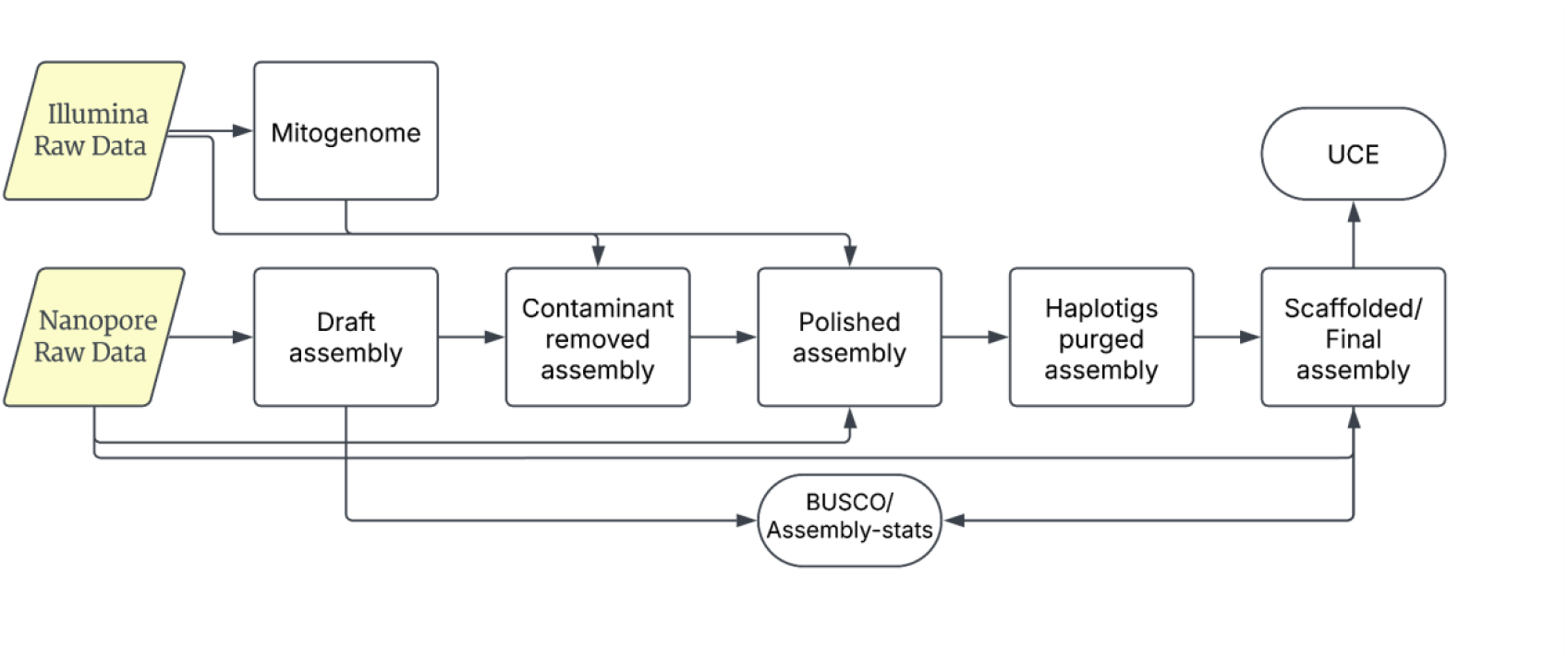
Schematic representation of the methodology followed to assemble the genomes.

### 2.6 Repeat Masking and Genome Annotations

We used RepeatModeler v2.0.2 (Flynn et al. 2020) within the Dfam TETools container v1.88 (Dfam Consortium 2023) with -LTRStruct to create a species-specific repeat element library. Species-specific libraries were merged with existing repeat libraries sourced from Dfam 0th partition (Storer et al. 2021) and RepBase RepeatMasker libraries (v20181026) (Jurka et al. 2005; Bao, Kojima, and Kohany 2015). The resulting combined library was then used to identify and soft mask (−xsmall) the repeat elements using RepeatMasker v4.1.2-p1 (Smit, Hubley and Green 2013). We then utilized Braker v3.0.3 (Stanke et al. 2008, 2006; Brůna et al. 2021) to predict the gene models. BRAKER predictions are based on the successive training of GeneMark-EP+ and AUGUSTUS with extrinsic evidence of homologous protein sequences. We employed the ProtHint pipeline within BRAKER and trained AUGUSTUS using vertebrate amino acid sequences from Vertebrata_OrthoDB_10 (Brůna, Lomsadze, and Borodovsky 2020; Lomsadze et al. 2005; Iwata and Gotoh 2012; Gotoh, Morita, and Nelson 2014; Buchfink, Xie, and Huson 2015). We first converted the BRAKER gtf to gff using agat v1.4.0 (Dainat et al. 2024) and sanitized the gff3 with gfftk v24.2.4 (https://github.com/nextgenusfs/gfftk). We then generated functional annotations using InterProScan v5.66 -98.0 (Jones et al. 2014) and eggNOG-mapper v2 (with eggNOG v5 database) (Cantalapiedra et al. 2021; Huerta-Cepas et al. 2019; Buchfink, Reuter, and Drost 2021), and the resulting functional annotations were integrated using containerized funannote v1.8.15 (Palmer 2023) and summarized with gfftk.

### 2.7 Global Avian Assembly Comparison

We used NCBI’s genome page (https://www.ncbi.nlm.nih.gov/datasets/genome/) to search ‘aves’ as a keyword and the Datasets v15.19.0 (O’Leary et al. 2024) command line tool to retrieve the information on all available avian genomes as of March 06, 2025. We retained only the latest genome for a species when more than one genome was available. We compared the statistics between existing avian assemblies and genomes generated in this study, such as contig N50, scaffold N50, and the number of contigs and scaffolds. Because the scaffold numbers spanned several orders of magnitude, each count was log10 transformed and plotted as a histogram in RStudio with R 4.4. (Posit team 2024). The median scaffold counts were calculated to compare the assembly contiguity between the existing and newly generated assemblies, and a two-sided Wilcoxon rank-sum test was performed using the ‘wilcox.test’ function in R.

## 3 Results

### 3.1 Raw Sequencing Output

Long read sequencing yielded 19.7 million reads with an average of 2.8 million reads per sample and an average read N50 of 12.89 Kbp across samples. 97.1% of the total long reads passed the QC. Additionally, Illumina sequencing resulted in 693.96 million reads with an average of 99.13 million reads per sample. Adapter trimming and QC dropped only 0.02% of the short reads (Appendix S1). GenomeScope2 estimated the genome size to be between 987 Mbp (*Machlolophus aplonotus*) and 1.12 Gbp (*Harpactes fasciatus*) with a maximum heterozygosity of 0.83% for *Geokichla citrina*. See supplementary Figure S1 for the individual GenomeScope2 profile.

### 3.2 Nuclear Genome Assembly

Assembled genome size ranged between 1.06 Gbp for *Alcippie poioicephla* and 1.13 Gbp for *Nyctyornis athertoni*. The least fragmented assembly was of *Hypothymis azurea* with 1775 scaffolds, and the most fragmented was *Myophonus horsfieldii* with 3886 scaffolds. Detailed genome assembly statistics are given in Table 1. In general, haplotig purging and polishing reduced the number of contigs from the initial draft assembly, and scaffolding with ntLink increased the contiguity. Contig N50 ranged between 653 Kbp (*Harpactes fasciatus)* and 2.10 Mbp (*Geokichla citrina)*, whereas the scaffold N50 ranged between 655 Kbp (*Harpactes fasciatus)* and 2.12 Mbp (*Geokichla citrina)*, with the largest scaffold size of 9.7 Mbp. Notably, Scaffold L50 for *Geokichla citrina* was 145, whereas that of *Harpactes fasciatus* was 512. Compleasm revealed that all the assemblies contained a high proportion of avian BUSCOs (n = 8337), with completeness exceeding 97% for all assemblies. Among the assemblies, *Machlolophus aplonotus* exhibited the highest BUSCO completeness (98.90%), followed by *Hypothymis azurea* (98.80%), *Geokichla citrina* (98.70%), *Myophonus horsfieldii* (98.50%), and *Alcippe poioicephala* (98.30%). *Harpactes fasciatus* and *Nyctyornis athertoni* assemblies showed slightly lower completeness, with 97.60% of complete BUSCOs recovered. Harvesting for UCEs resulted in a minimum of 4799 loci recovered across assemblies. Among the assemblies, we recovered the highest number of loci from *Hypothymis azurea* (4890), followed by *Myphonus horsfieldii* (4867), *Nyctyornis athertoni* (4859), *Harpactes fasciatus* (4856), *Machlolophus aplonotus* (4848), and *Geokichla citrina* (4825). The least number of loci was recovered from *Alcippe poiociephala* (4799).

**Table 1:**
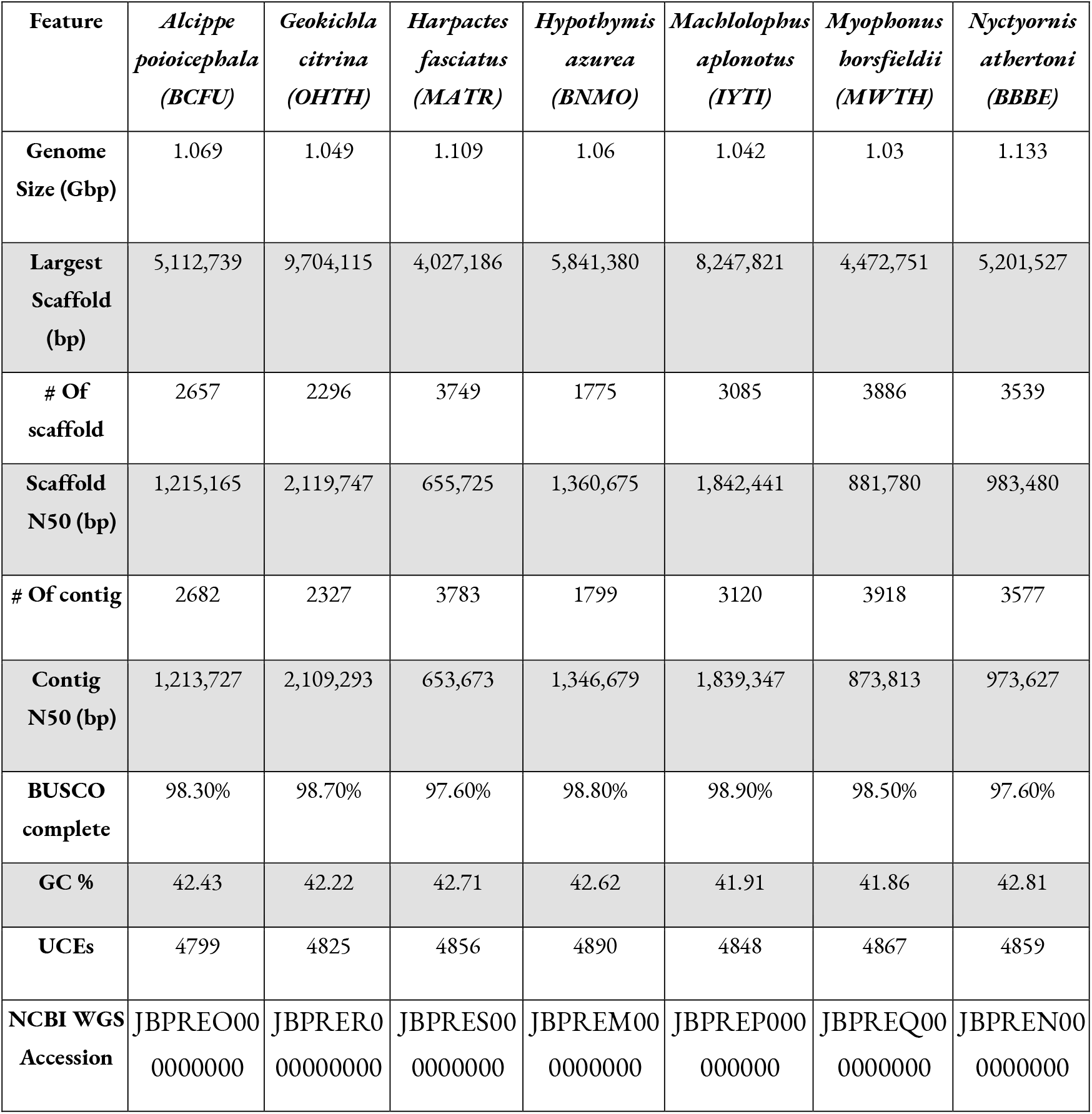
Genome assembly contiguity and completeness characteristics for the seven family assemblies.

### 3.3 Mitochondrial Genome Assembly

The final mitochondrial assembly size ranged from 14,066 bp to 17,888 bp. Our mitogenomes comprise 13 (IYTI, MWTH, OHTH, BCFU) or 12 protein-coding genes (BBBE, MATR, BNMO) with 20-22 tRNAs and two rRNAs. *Harpactes fasciatus* had the least GC content at 43%, and *Machlolophus aplonotus* had the most at 49%. Most of the protein-coding genes were on the heavy chain of the mitogenome across species, with a one or both D-loop for the circularized mitogenomes. See supplementary Table S2 for the individual mitogenome statistics.

### 3.4 Repeat Masking and Genome Annotations

RepeatMasker identified interspersed repeats ranging from 8.44% in *Myophonus horsfieldii* to 11.07% in *Nyctyornis athertoni*. Most of the repetitive elements were retroelements, followed by simple and unclassified repeats, while satellites and DNA transposons comprised a smaller proportion across assemblies (Figure 3). Notably, *Geokichla citrina* exhibited a comparatively higher proportion of DNA transposons (0.78% v/s <0.13%) than the other assemblies (Appendix S2). Gene prediction using the BRAKER pipeline resulted in an average of 35,551 putative gene models per genome, with an average of 37,605 mRNA transcripts. *Geokichla citrina* had the highest number of predicted gene models (42,048), while *Nyctyornis athertoni* had the fewest (29,277). On average, 9,619 gene models per genome were assigned functional annotations (See supplementary Figure S4).

**Figure 3:**
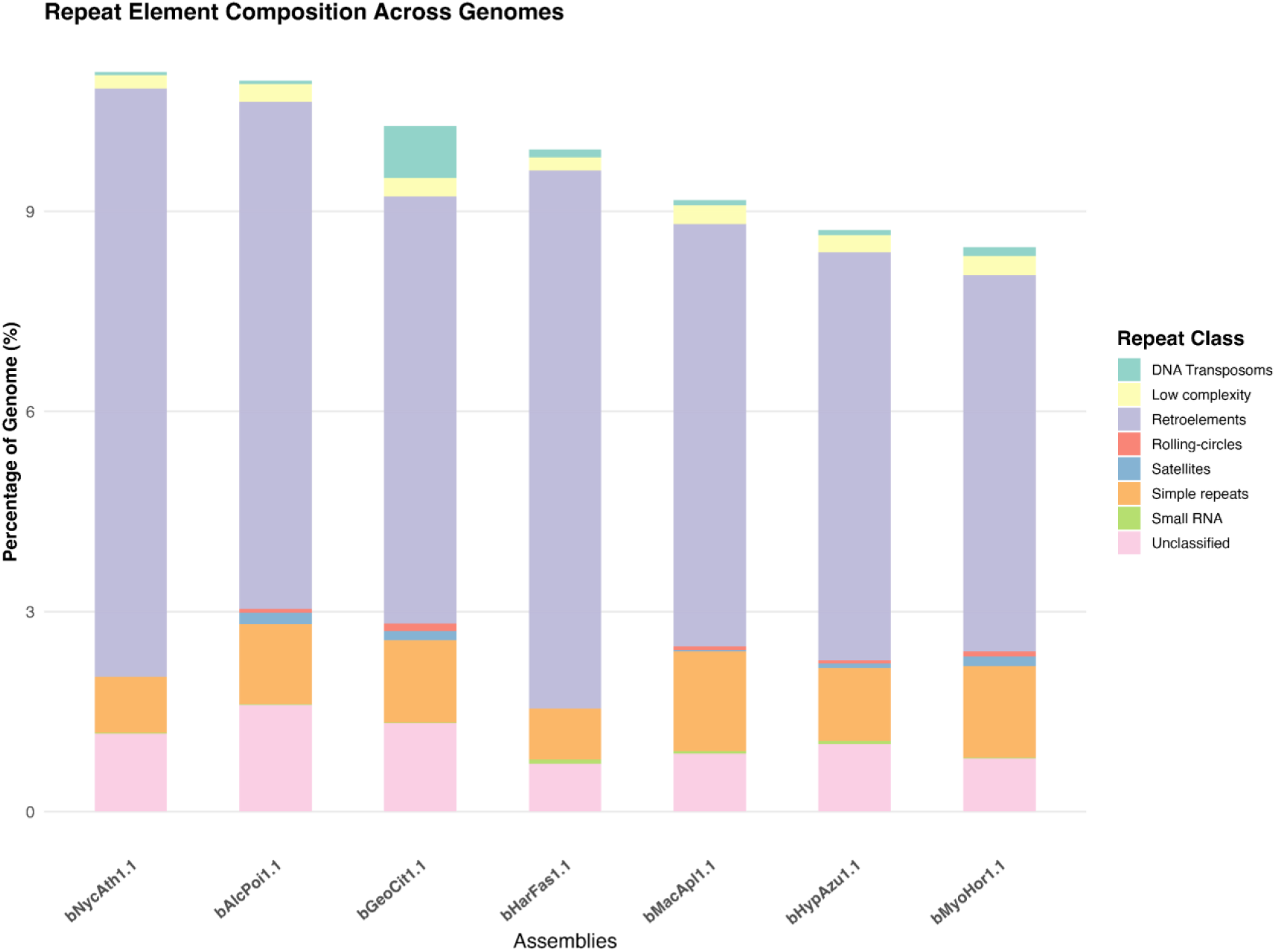
Repeat element composition across seven assemblies. Stacked bar plots show the percentage of the genome occupied by major repeat classes (y-axis) across each assembly (x-axis), including unclassified repeats, small RNA, simple repeats, satellites, rolling-circles, retroelements, low-complexity, and DNA transposons as identified by RepeatMasker. Retroelements account for the largest fraction of repeats across all assemblies, followed by unclassified and simple repeats. Notably, the bGeoCit1.1 displays a higher proportion of DNA transposons compared to others.

### 3.5 Global Avian Assembly Comparison

Our search for the available bird genomes on NCBI yielded 2365 genomes. Taxonomic discrepancies between NCBI and BirdLife International resulted in a mismatch of 90 species, of which only 67 species could be resolved by manual assignment (Appendix S3). After filtering to remove subspecies designations and to retain only the latest assembly for each species, we were left with 1467 genomes representing 1467 species for downstream comparison. Our comparison of genomes generated in this study via a hybrid ONT-Illumina approach with existing assemblies exhibits improved contiguity relative to the existing avian assemblies. The median scaffold count for the newly generated assemblies is 3,085, substantially lower than the 28,877 scaffolds of the existing genome assemblies for birds (Wilcoxon rank-sum W = 1,697, p = 0.00226) – indicating fewer, larger scaffolds and thus achieving higher overall assembly contiguity (Figure 4A). Furthermore, the scatterplot of scaffold N50 versus the contig N50 confirms this enhancement of the assemblies, as all seven species’ genomes cluster in the upper-right quadrant (Figure 4B), indicating the improved contiguity relative to the bulk of the existing bird genomes.

**Figure 4:**
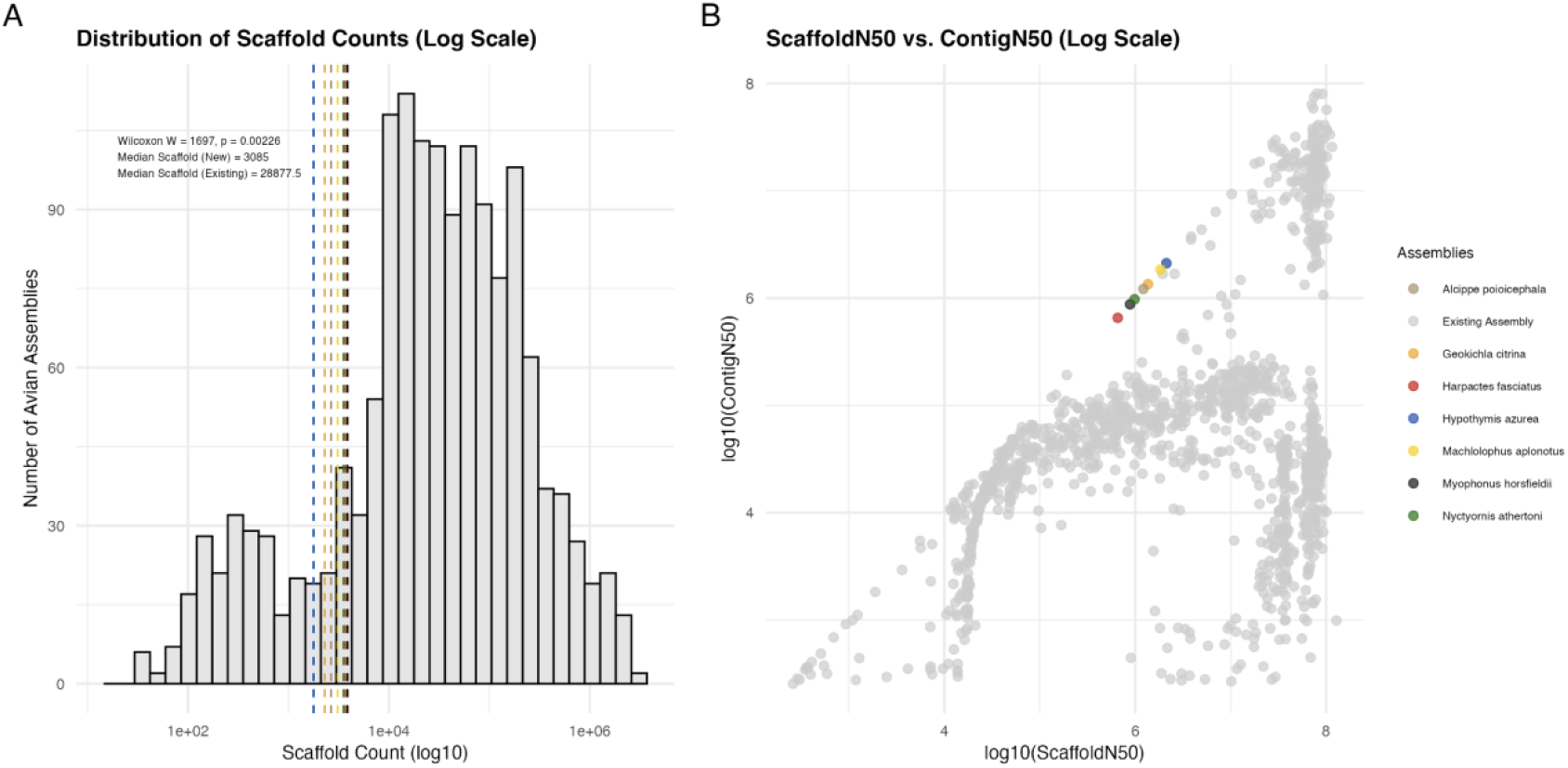
Assembly contiguity for the assemblies generated in this study versus the existing avian genomes. A) Histogram of scaffold counts on a log10 scale (grey bars), overlaid with dashed vertical lines representing the median scaffold count of newly generated (colored) assemblies. The new assemblies exhibit a significantly lower median scaffold (3,085 vs. 28,877; Wilcoxon W = 1,697, p = 0.00226), reflecting larger, more contiguous assemblies. B) Scatterplot of log10 (Scaffold N50) against log10 (Contig N50); existing genomes are in light grey and newly generated assemblies are in color. All new assemblies cluster in the upper-right quadrant, demonstrating the higher contiguity compared to the existing assemblies in NCBI.

## 4 Discussion

In recent years, there has been a surge in generating reference genomes targeting biodiversity studies (Formenti et al. 2022), and generating high-quality genomic resources for organisms originating from biodiverse rich, yet historically under-sampled regions, is critical for understanding avian evolution and conservation genomics. By generating high-quality genomes for seven species belonging to seven families of birds from the Western Ghats of India, our study adds taxonomic breadth to the public database of genomic resources.

Our final genome assembly sizes ranged between 1.06 Gbp and 1.13 Gbp, well within the expected genome length for birds (∼1.0 Gbp to ∼1.2 Gbp) (Kapusta, Suh, and Feschotte 2017) and similar to previously reported avian genomes (Feng et al. 2020). Across species, scaffold N50 values ranged from 0.66 Mbp to 2.12 Mbp (Table 1). These figures compare favorably with the median scaffold N50 of existing avian genomes assembled with short-read data alone. Genome assemblies reported here are more contiguous than the bulk of the existing assemblies (Figure 4), and BUSCO scores of >97% for all assemblies and >4800 recovered UCE loci indicate a high proportion of completeness of the assemblies. Our final reported mitogenomes are circular and similar in size to those previously reported for passerines and other birds (Lan et al. 2024; Lu et al. 2019; Feng et al. 2020). The total number of protein-coding genes, rRNAs, and tRNAs is comparable to the expected number of elements in a vertebrate mitogenome (Formenti et al. 2021), making them suitable to use in the clade-wide mitogenome studies.

Our assembly comparison results highlight how much variation there still is in the publicly available bird genomes. While the number of assemblies has grown potentially driven by the global sequencing initiatives such as the Bird 10K project (Zhang et al. 2015), the Earth Bio Genome Project (Lewin et al. 2018), the Darwin Tree of Life Project (Darwin Tree of Life Project Consortium 2022), and the European Reference Genome Atlas (Mc Cartney et al. 2024) which aims to generate reference assemblies for all the existing vertebrate species, many are still quite fragmented. In contrast, the assemblies presented here show much higher contiguity (Figure 4), making them useful for looking at the large-scale patterns across clades. The global analysis of tetrapods (Linck and Cadena 2024), amphibians and reptiles (Carneiro et al. 2025) found that the Global South is substantially underrepresented in the genomic resources and available reference genomes, including biodiverse, rich tropical regions, such as Southeast Asia, and much of the Indian subcontinent. Although a few genome assemblies are associated with species found in the Indian Peninsula, most are derived from taxa with wide geographic distribution, such as the Great Egret (*Ardea alba*; GCA_045788385.1), Red-necked Falcon (*Falco chicquera*; GCA_034781875.1), and Black-crowned Night Heron (*Nycticorax nycticorax*; GCA_023375905.1). Some genomes are based on the introduced populations, such as the Red-whiskered Bulbul (*Pycnonotus jocosus*; GCA_013400435.1) or from captive, including zoo-sourced samples [for example, White-rumped Munia (*Lonchura striata*; GCA_046129705.1), Indian Peafowl (*Pavo cristatus*; GCA_045791835.1, GCA_005519975.1, GCA_965225075.1)] and private breeders [for example, Lady Amherst’s pheasant (*Chrysolophus amherstiae*; GCA_036784685.1)]. In contrast, a handful of genome assemblies are assembled from wild-caught birds originating from India. Notably, only a couple of assemblies from the Western Ghats (long read technologies; Pawar et al. 2025; Vinay et al. 2025) and an additional couple of assemblies from the other regions of India (short read technologies; Kumar et al. 2023; Mondal et al. 2023).

The high-quality, annotated genome assemblies presented here for seven passerines will be valuable for understanding the population structure, biogeographic patterns, and colonization of birds in peninsular India. The genome assemblies presented here substantially improve avian genomes’ taxonomic and geographic coverage in the Global South, with assembly metrics adequate for most molecular-ecology applications. While PacBio HiFi reads, coupled with hi-C data for scaffolding, would help us achieve the chromosome-level and inch towards the Vertebrate Genome Project recommendation metrics for a reference genome (Rhie et al. 2021), it is not feasible to collect fresh tissue samples due to recent changes in the legal provisions in obtaining permits to capture the birds (Shanker et al. 2023; Indian Environmental Portal 2025). Given the legal difficulties in obtaining fresh samples and fieldwork (Madhusudan et al. 2006), the genome resources from wild-caught birds from the underrepresented regions will be far more valuable to scientific communities. We hope these resources provide a foundation for comparative and population genomic studies of the South Asian birds, and they will be of interest to a broader audience in evolutionary biology and contribute to the ever-growing repository of avian genome assemblies.

## Author Contributions

VKL, NG, and VVR conceptualised the research, VKL and NG designed and executed the study. NG, AW, and CA collected samples and conducted laboratory work. VKL and NG performed the study and analysed the data. VKL generated the figures and wrote the draft manuscript. All authors reviewed and edited the manuscript. VVR raised funds and administered the project.

## Acknowledgments

This study was funded by Rohini Nilekani Philanthropies Foundation and the Indian Institute of Science Education and Research (IISER) Tirupati internal funds. We thank Kerala and Tamil Nadu State Forest Departments for the field support and permits. We thank the IISER Tirupati IT department for their computational support during this study. We are grateful to Murugavel, Kovai Thambi, Kamraj, Eswaran, Shivam Shinde, Viral Joshi, and Aishwarya Bhandari for their help during the field work in Kerala. We thank Vishwa Jagati and Gayathri R for helping with lab work.

## Data accessibility statement

All raw reads are available in NCBI under BioProject PRJNA1290491 and Sequence Read Archive (SRA) accession SRR34505061-SRR34505067 (for ONT reads) and SRR34530187-SRR34530193 (for Illumina reads). Genome assemblies are available at NCBI under WGS accessions as listed in Table 1 for the respective species. Predicted and annotated genes are deposited in the Open Science Framework (OSF) osf.io/uwa53, and the scripts are deposited in GitHub (https://github.com/stachyris/PeninsularBirdAssemblies).

## Benefit sharing statement

Benefits Generated: Benefits from this research accrue from the sharing of our data and results on public databases as described above.

